# Robust expression variability testing reveals heterogeneous T cell responses

**DOI:** 10.1101/237214

**Authors:** Nils Eling, Arianne C. Richard, Sylvia Richardson, John C. Marioni, Catalina A. Vallejos

## Abstract

Cell-to-cell transcriptional variability in otherwise homogeneous cell populations plays a crucial role in tissue function and development. Single-cell RNA sequencing can characterise this variability in a transcriptome-wide manner. However, technical variation and the confounding between variability and mean expression estimates hinders meaningful comparison of expression variability between cell populations. To address this problem, we introduce a novel analysis approach that extends the BASiCS statistical framework to derive a residual measure of variability that is not confounded by mean expression. Moreover, we introduce a new and robust procedure for quantifying technical noise in experiments where technical spike-in molecules are not available. We illustrate how our method provides biological insight into the dynamics of cell-to-cell expression variability, highlighting a synchronisation of the translational machinery in immune cells upon activation. Additionally, our approach identifies new patterns of variability across CD4^+^ T cell differentiation.

## Introduction

Heterogeneity in gene expression within a population of single cells can arise from a variety of factors. Structural differences in gene expression within a cell population can reflect the presence of sub-populations of functionally different cell types (Zeisel et al., 2015; Paul et al., 2015). Alternatively, in a seemingly homogeneous population of cells, so-called unstructured expression heterogeneity can be linked to intrinsic or extrinsic noise (Elowitz et al., 2002). Changes in physiological cell states (such as cell cycle, metabolism, abundance of transcriptional/translational machinery and growth rate) represents extrinsic noise, which has been found to influence expression variability within cell populations (Keren et al., 2015; Buettner et al., 2015; Zeng et al., 2017). Intrinsic noise can be linked to epigenetic diversity (Smallwood et al., 2014), chromatin rearrangements (Buenrostro et al., 2015), as well as the genomic content of single genes, such as the presence of TATA-box motifs and the abundance of nucleosomes around the transcriptional start site (Hornung et al., 2012).

Single-cell RNA sequencing (scRNAseq) generates transcriptional profiles of single cells, allowing the study of cell-to-cell heterogeneity on a transcriptome-wide (Grün et al., 2014) and single gene level (Goolam et al., 2016). Consequently, this technique can be used to study unstructured cell-to-cell variation in gene expression within and between homogeneous cell populations (i.e. where no distinct cell sub-types are present). Increasing evidence suggests that this heterogeneity plays an important role in normal development (Chang et al., 2008) and that control of expression noise is important for tissue function (Bahar Halpern et al., 2015). For instance, molecular noise was shown to increase before cells commit to lineages during differentiation (Mojtahedi et al., 2016), while the opposite is observed once an irreversible cell state is reached (Richard et al., 2016). A similar pattern occurs during gastrulation, where expression noise is high in the uncommitted inner cell mass compared to the committed epiblast and where an increase in heterogeneity is observed when cells exit the pluripotent state and form the uncommitted epiblast (Mohammed et al., 2017).

Motivated by scRNAseq, recent studies have extended traditional differential expression analyses to explore more general patterns that characterise differences between cell populations (e.g. Korthauer et al., 2016). In particular, BASiCS (Vallejos et al., 2015, 2016) introduced a probabilistic tool to assess differences in cell-to-cell heterogeneity between two or more cell populations. To meaningfully assess these changes, BASiCS separates biological noise from technical variability — a characteristic feature of scRNAseq datasets (Brennecke et al., 2013) — by borrowing information from synthetic RNA spike-in molecules.

The differential test implemented in BASiCS has led to, for example, novel insights in the context of immune activation and ageing (Martinez-Jimenez et al., 2017). However, a major challenge remains unresolved. Biological noise is negatively correlated with protein abundance (Bar-Even et al., 2006; Newman et al., 2006; Taniguchi et al., 2010) or mean RNA expression (Brennecke et al., 2013; Antolović et al., 2017). Therefore, changes in variability between two populations of single cells are confounded by changes in mean expression.

This article extends the statistical model implemented in BASiCS by deriving a residual measure of cell-to-cell transcriptional variability that is not confounded by mean expression. In addition, we address experimental designs where spike-in sequences are not available by exploiting concepts from measurement error models.

Using our method, we identify a synchronisation of the translation machinery in CD4^+^ T cells upon early immune activation as well as an increased variability in the expression of genes related to CD4^+^ T cell immunological function. Furthermore, we detect evidence of early cell fate commitment of CD4^+^ T cells during malaria infection characterized by a decrease in *Tbx21* expression heterogeneity and a rapid collapse of global transcriptional variability after infection. These results lead to new insights into transcriptional dynamics during immune activation and differentiation and propose an earlier time point of cell fate commitment than previously anticipated.

## Results

### Addressing the mean confounding effect for differential variability testing

Unlike bulk RNA sequencing, scRNAseq provides information about cell-to-cell expression heterogeneity within a population of cells. Past works have used a variety of measures to quantify this heterogeneity. Among others, this includes the coefficient of variation (CV, Brennecke et al., 2013) and entropy measures (Richard et al., 2016). As in Vallejos et al. (2015, 2016), we focus on *over-dispersion* as a proxy for transcriptional heterogeneity. This is defined by the excess of variability that is observed with respect to what would be predicted by Poisson sampling noise, after accounting for technical variation.

The aforementioned measures of variability can be used to identify genes whose transcriptional heterogeneity differs between groups of cells (defined by experimental conditions or cell types). However, the strong relationship that is typically observed between mean and variability estimates (e.g. Brennecke et al., 2013) can hinder the interpretation of these results.

A simple solution to avoid this confounding is to restrict the assessment of differential variability to those genes with equal mean expression across populations (see **Fig. 1A**, also Martinez-Jimenez et al., 2017). However, this is sub-optimal, particularly when a large number of genes are differentially expressed between the populations. For example, reactive genes that change in mean expression upon changing conditions (e.g. transcription factors) are excluded from differential variability testing. An alternative approach is to directly adjust variability measures to remove this confounding. For example, Kolodziejczyk et al. (2015) computed the distance between the squared CV to a rolling median along expression levels — referred to as the DM method. However, the DM method does not allow statistical testing of differences at a gene level and does not take into account technical variation.

**Figure 1:**
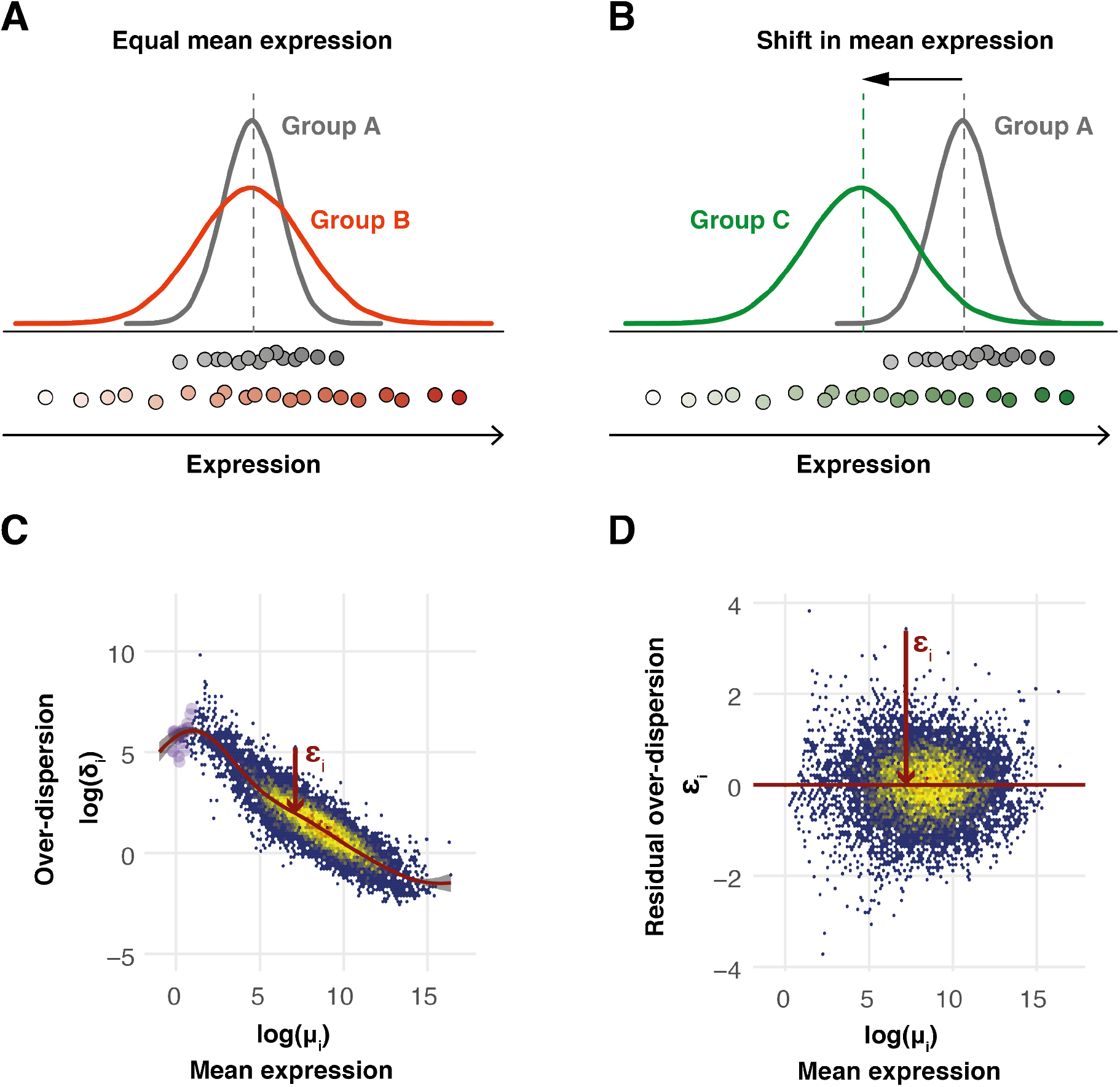
Avoiding the mean confounding effect when quantifying expression variability in scRNAseq data. (A)-(B) Illustration of changes in expression variability for a single gene between two cell populations without (A) and with (B) changes in mean expression. (C)-(D) BASiCS parameters were estimated using the dataset introduced by Antolović et al. (2017). Genes that are not expressed in at least 2 cells are indicated by purple points. The red line shows the regression trend between over-dispersion and mean expression. Residual overdispersion is indicated for one gene by a red arrow. (C) BASiCS’s estimates of gene-specific over-dispersion *δ_i_* are confounded by mean expression estimates *μ_i_* in the context of scRNAseq datasets. (D) After performing the regression between *δ_i_* and *μ_i_*, residual over-dispersion estimates do not correlate with mean expression.

Our method solves this problem by extending the statistical model implemented in BASiCS (Vallejos et al., 2015, 2016). We define a measure of *residual over-dispersion* — which is not correlated with mean expression — to meaningfully assess changes in transcriptional heterogeneity when genes exhibit shifts in mean expression (see **Fig. 1B**). We infer a regression trend between gene-specific mean (*μ_i_*) and over-dispersion parameters (*δ_i_*), by introducing a joint informative prior for these parameters (see **Methods**). A gene-specific *residual over-dispersion* parameter *ϵ_i_* describes departures from this trend (see **Fig. 1C**). Positive values of *ϵ_i_* indicate that a gene exhibits more variation than expected relative to genes with similar expression levels. Similarly, negative values of *ϵ_i_* suggest less variation than expected. Importantly, as shown in **Fig. 1D**, these residual over-dispersion parameters are not confounded by mean expression. Thus, *ϵ_i_* can be used to meaningfully assess changes in transcriptional heterogeneity, regardless of mean expression differences.

### The informative prior stabilizes parameter estimation

To study the effect of the joint informative prior described above, we applied our method to a variety of scRNAseq datasets. Each dataset is unique in its composition, covering a range of different cell types and experimental protocols (see **Supplementary Note S3** and **Table S1**). Qualitatively, the trend between the mean *μ_i_* and the over-dispersion *δ_i_* varies substantially across datasets, justifying the choice of a flexible semi-parametric approach for capturing this trend (see **Methods** and **Fig. S1**). Moreover, in line with Love et al. (2014), the introduction of a joint prior specification for (*μ_i_,δ_i_*)′ regularises parameter estimation by shrinking the posterior estimates for *μ_i_* and *δ_i_* towards the estimated trend. The strength of this shrinkage is also dataset-specific and has the most impact for lowly-expressed genes where measurement error is greatest. Nevertheless, in all cases, we observe that residual over-dispersion parameters *ϵ_i_* are not confounded by mean expression, nor by the percentage of zero counts per gene (see **Fig. S1**).

Subsequently, we ask whether or not the shrinkage described above improves posterior inference. We assess estimation performance across different sample sizes by comparing posterior medians estimated by the regression and the non-regression models against a *pseudo* ground truth (**Fig. 2, Fig. S2** and **Supplementary Note S5.1**). The latter is derived through a sub-sampling strategy in which empirical estimates obtained on the basis of a large sample size are used as a proxy for the true values of *μ*_i_, *δ_i_* and *ϵ_i_*. This is based on a subset of the data described in Zeisel et al. (2015), containing 939 CA1 pyramidal neurons. Subsequently, we use random sub-sampling to generate datasets containing *n* cells (50 ≤ *n* ≤ 500) and calculate the associated posterior estimates. While the shrinkage effect does not alter posterior inference for mean expression parameters *μ_i_* (**Fig. 2A**), a more prominent effect is observed for over-dispersion parameters *δ_i_* (**Fig. 2B**). The latter is emphasized when the data are less informative, such as for low sample sizes, where it is more likely to observe zero total counts (across all cells) for lowly expressed genes. In those cases, shrinkage towards the trend avoids artificially low estimates for *δ*_i_. Overall, in terms of the residual over-dispersion parameters *ϵ_i_*, the property yields statistically unbiased estimates (**Fig. 2C**). However, a small over-estimation of *ϵ_i_* can be observed for lowly expressed genes when the sample size is small (this bias is four times smaller than the default threshold for differential variability testing, Ψ_0_ > 0.41).

**Figure 2:**
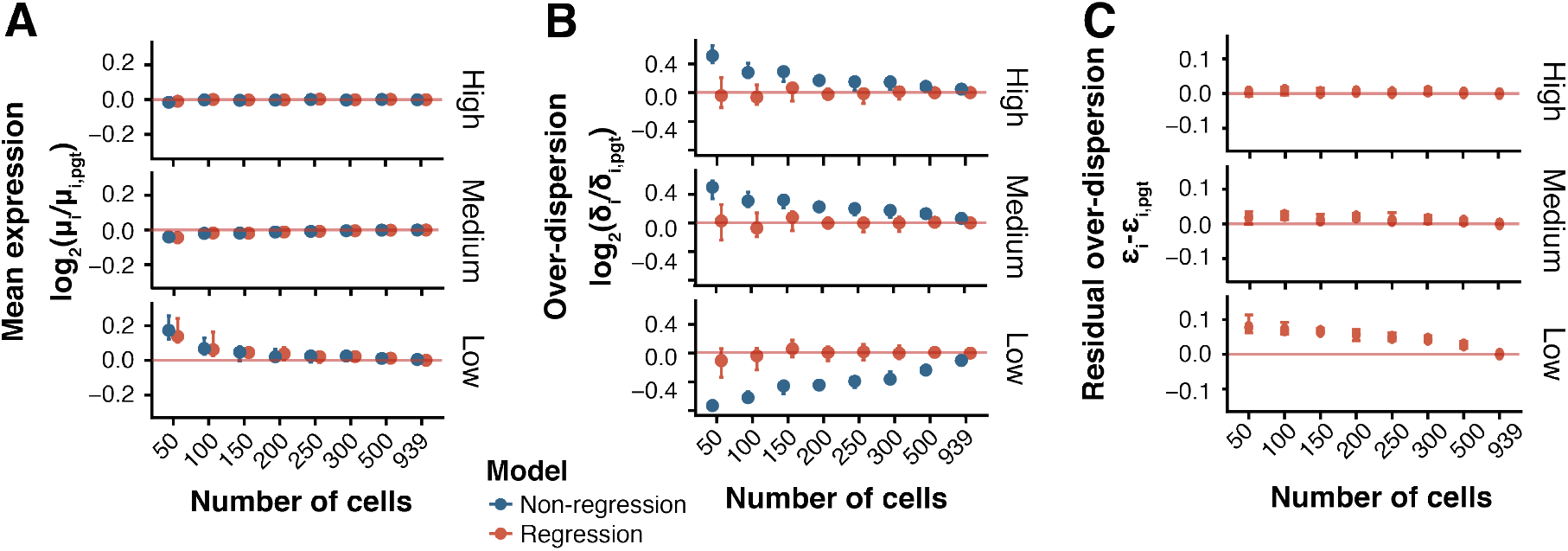
Parameter estimation for varying cell population size. The regression (orange) and non-regression (blue) model was used to estimate model parameters for lowly (lower panels), medium (mid panels) and highly (upper panels) expressed genes across populations of 50-500 cells. Parameters estimated on the full population of 939 cells were used as *pseudo* ground truth (pgt). Median log_2_ fold changes (*μ_i_* and *δ_i_*) and median distances (*ϵ_i_*) between the estimates and the pgt were computed. The median value across 10 replicates is plotted with error bars indicating the range across all repetitions (see **Supplementary Note S5.1**). (A) Distribution of log_2_ fold changes in mean expression *μ_i_* estimates. Default threshold for differential mean expression testing: 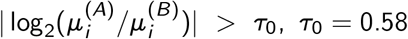. (B) Distribution of log_2_ fold changes in over-dispersion *δ_i_* estimates. Default threshold for differential over-dispersion testing: 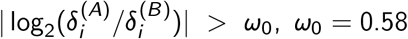. (C) Distribution of absolute distances for residual over-dispersion *ϵ_i_* estimates. Default threshold for differential mean expression testing: 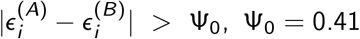.

### Inferring technical variability without spike-in genes

Another critical aspect to take into account when inferring transcriptional variability based on scRNAseq datasets is technical variation (Brennecke et al., 2013). BASiCS (and its extension described above) achieves this by exploiting a set of synthetic RNA spike-in molecules (e.g. the set of 92 ERCC molecules developed by Jiang et al., 2011) as a *gold standard* to aid normalisation and to quantify technical artefacts. However, while the addition of spike-in genes prior to sequencing is theoretically appealing (Lun et al., 2017), several practical limitations can preclude their utility in practice (Vallejos et al., 2017).

Given this, we extend BASiCS so that it can handle datasets without spike-in genes by exploiting principles of measurement error models where — in the absence of gold standard features — technical variation is quantified through *replication* (Carroll, 1998). This is based on experimental designs where cells from a population are randomly allocated to multiple independent experimental replicates (here referred to as *batches).* In such an experimental designs, the no-spikes implementation of BASiCS assumes that biological effects are shared across batches and that technical variation will be reflected by spurious differences. As shown in **Fig. S3E-F**, posterior inference under the no-spikes BASiCS model closely matches the original implementation for datasets where spike-ins and batches are available. Technical details about the no-spikes implementation of BASiCS are discussed in **Supplementary Note S4**.

### Expression variability dynamics during immune activation and differentiation

Here, we illustrate how our method assesses changes in expression variability using CD4^+^ T cells as model system. For all datasets, pre-processing steps are described in **Supplementary Note S3**.

### Testing variability changes in immune response gene expression

To identify gene expression changes during early T cell activation, we compared CD4^+^ T cells before (naive) and after (active) 3h of stimulation (Martinez-Jimenez et al., 2017). Previously, Martinez-Jimenez et al. (2017) observed that activated CD4^+^ T cells synchronise their expression upon activation, solely focusing on genes with no changes in mean expression. This represents only ~2,000 out of the full set of ~10,000 expressed genes. In contrast, testing changes in variability through residual over-dispersion allows us to include the large set of genes that are up-regulated upon immune activation. Importantly, these include immune-response genes and critical drivers for CD4^+^ T cell functionality.

Our model classifies genes into four categories based on their expression dynamics: down-regulated upon activation with (i) lower and (ii) higher variability, and up-regulated with (iii) lower and (iv) higher variability (**Fig. 3A** and **Supplementary Note S5.2**).

**Figure 3:**
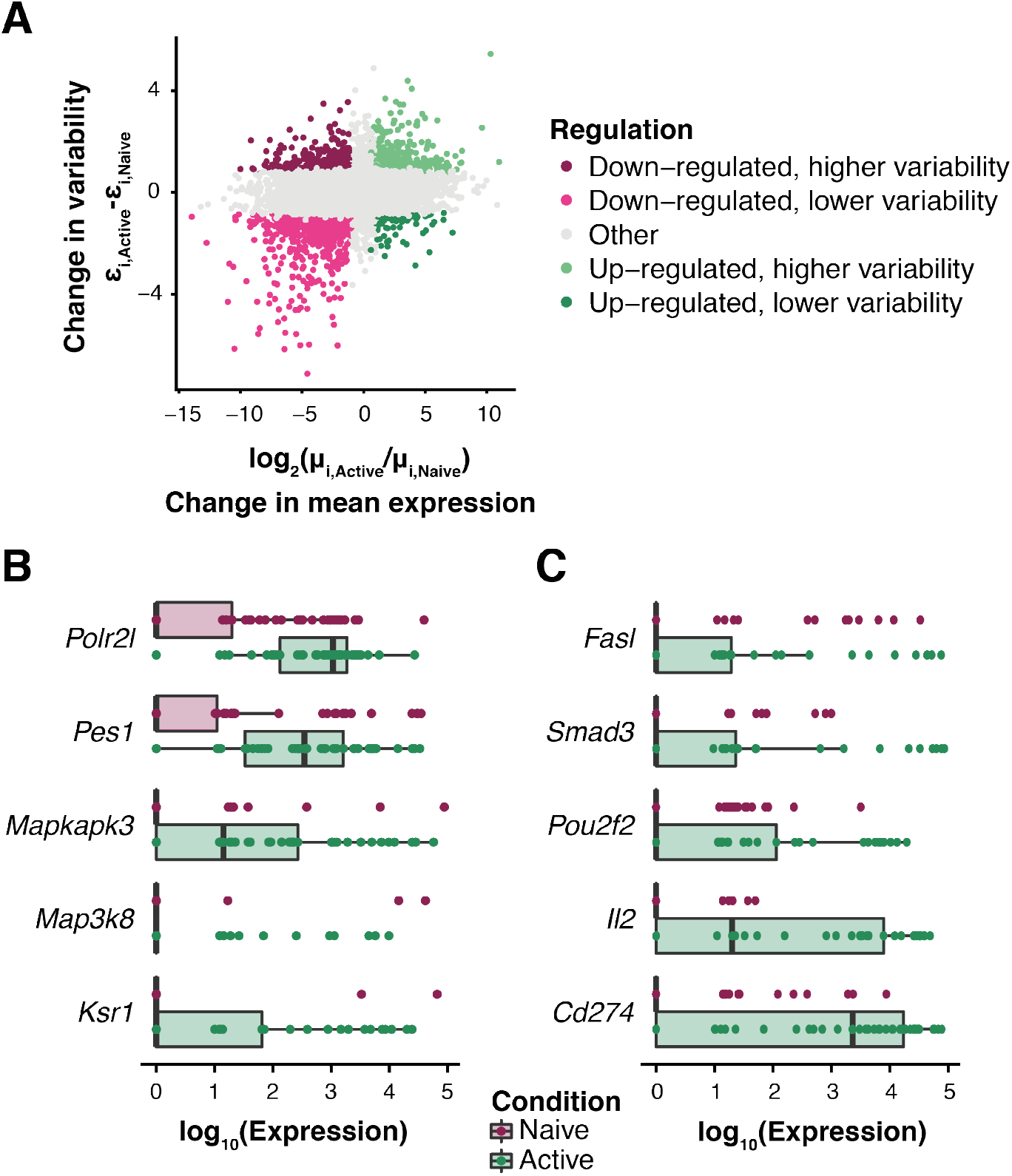
Changes in expression variability during early immune activation in CD4^+^ T cells. Differential testing was performed between naive and activated murine CD4^+^ T cells testing a log_2_ fold change in mean expression > 1 and an absolute distance between residual over-dispersion Ψ_0_ > 0.41 while controlling the expected false discovery rate at 10% (see **Supplementary Note S5.2**). (A)For each gene, the distance in residual over-dispersion estimates (Active - Naive) is plotted versus the log_2_ fold change in mean expression (Active/Naive). Genes with statistically significant changes in mean expression and variability are coloured based on their regulation (up/down-regulated, more/less variability). (B) Denoised expression across the naive (purple) and active (green) CD4^+^ T cell population is visualized for representative genes that increase in mean expression and decrease in expression variability upon immune activation. Each dot represents a single cell. (C) Distribution of denoised expression counts for sample genes that increase in mean expression as well as expression variability. Each dot represents a single cell.

Genes with up-regulated expression upon activation and decreased expression variability encode components of the MAP-kinase signaling pathways (e.g. *Mapkapk3, Map3k8, Ksr1*), RNA polymerase subunits (e.g. *Polr2l*), and ribosomal biogenesis and translation machinery components (e.g. *Pes1*) (**Fig. 3B**). These transcription factor-inducing signalling cascade and protein-building processes are required for naive T cells to rapidly enter a program of proliferation and effector molecule synthesis (Voll et al., 2000; Navarro and Cantrell, 2014). Therefore, rapid, uniform up-regulation of these transcripts would assist such processes. This observation also confirms previous findings that the translational machinery is tightly regulated during early immune activation (Martinez-Jimenez et al., 2017).

In contrast, immune-related genes with up-regulated expression and increased expression variability include the death-inducing and inhibitory transmembrane ligands Fas ligand (*Fasl*) and PD-L1 (*Cd274*), the regulatory transcription factor Smad3 (*Smad3*), and the TCR-induced transcription factor, Oct2 (*Pou2f2*). Additionally, we detect a heterogeneous up-regulation in the mRNA expression of the autocrine/paracrine growth factor IL-2 (*Il2*) upon immune activation (**Fig. 3C**). This is in line with previous reports of binary IL-2 expression within a population of activated T cells (Bucy et al., 1994), which has been suggested to be necessary for a scalable antigen response (Fuhrmann et al., 2016). Heterogeneity in expression of these genes suggests that, despite their uniform up-regulation of biosynthetic machinery, the T cells in this early activation culture represent a mixed population with varying degrees of activation and/or regulatory potential.

In summary, our approach allows us to extend the finding by Martinez-Jimenez et al. (2017), dissecting immune-response genes into two functional sets: (i) homogeneous up-regulation of translational machinery and (ii) heterogeneous up-regulation of immunoregulatory genes.

### Expression dynamics during *in vivo* CD4^+^ T cell differentiation

In contrast to the quick transcriptional switch that occurs within hours of naive T cell activation, transcriptional changes during cellular differentiation processes are more subtle and were found to be coupled with changes in variability prior to cell fate decisions (Richard et al., 2016; Mojtahedi et al., 2016). Here, we apply our method to study changes in expression variability during CD4^+^ T cell differentiation after malaria infection. In particular, we focus on samples collected 2, 4 and 7 days post-malaria infection, for which more than 50 cells are available (Lönnberg et al., 2017).

To study global changes in over-dispersion along the differentiation time course, we compare posterior estimates for the gene-specific parameter *δ_i_,* focusing on the 126 genes for which mean expression does not change (see **Fig. 4A** and **Supplementary Note S5.3**). This analysis suggests that the expression of these genes is most tightly regulated at day 4, when cells are in a highly proliferative state. Moreover, between day 4 and day 7, the cell population becomes more heterogeneous. This is in line with the emergence of differentiated Th1 and Tfh cells that was observed by Lönnberg et al. (2017).

**Figure 4:**
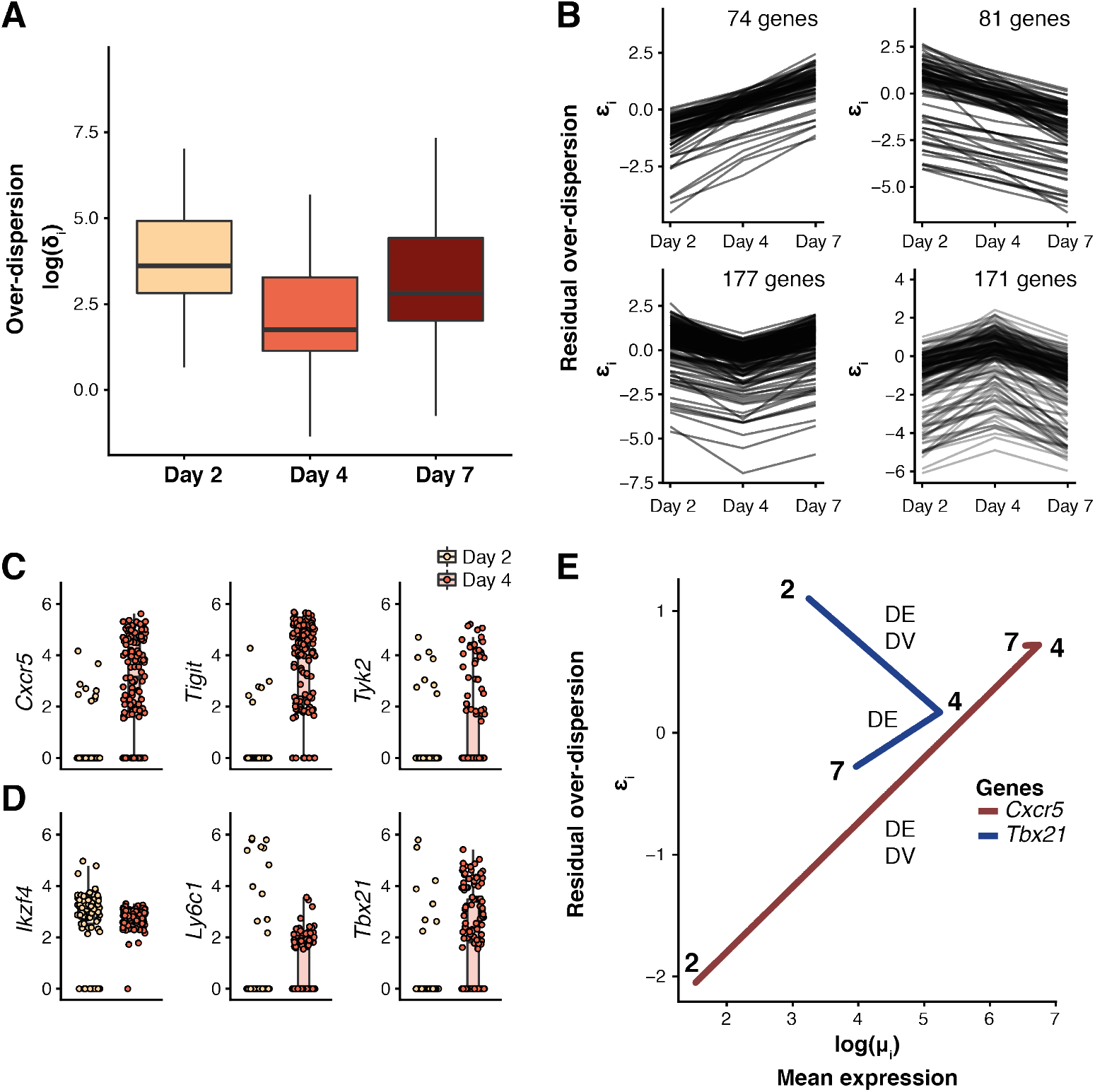
Dynamics of expression variability throughout CD4^+^ T cell differentiation. Analysis was performed on CD4^+^ T cells 2 days, 4 days and 7 days after *Plasmodium* infection. Changes in residual over-dispersion were tested using a threshold of Ψ_0_ > 0.41, EFDR=0.1 (see **Supplementary Note S5.3**) (A) Distribution of over-dispersion parameter estimates *δ_i_* for genes that don’t change in mean expression across the differentiation time course (log_2_ fold change in mean expression > 0, EFDR=0.1). (B) Residual over-dispersion parameter estimates *ϵ_i_* for genes with statistically significant change in expression variability between time points. Gene set size is indicated for each plot. (C) Denoised, log_10_-transformed expression across cell populations at day 2 (yellow) and day 4 (red) post infection is visualized for representative genes that increase in variability during differentiation. Each dot represents a single cell. (D) Distribution of denoised, log_10_-transformed expression counts for representative genes that decrease in expression variability during differentiation. Each dot represents a single cell. (E) Residual over-dispersion estimates *ϵ_i_* are plotted against mean expression *μ_i_* for *Tbx21* (blue) and *Cxcr5* (red) measured at day 2, day 4 and day 7 post-infection. Statistically significant changes in mean expression (DE) and variability (DV) are indicated for each comparison.

We next exploit the residual over-dispersion parameters to identify genes with a statistically significant change in variability (irrespective of changes in mean expression) between consecutive time points (see **Supplementary Note S5.3**). Separating these genes by whether their variability increases or decreases between time points revealed four different patterns (**Fig. 4B**). These include genes whose variability systematically increases (or decreases) as well as patterns where variability is highest (or lowest) at day 4.

In particular, we detect statistically significant changes in expression variability for a set of immune-related genes between day 2 and day 4 post-infection (**Fig. 4C**). For example, expression of *Cxcr5* which encodes the chemokine receptor that directs Tfh cells to the B cell follicles (Crotty, 2014), strongly increases in variability on day 4. This finding agrees with results from Lönnberg et al. (2017), where Tfh differentiation was observed to be transcriptionally detectable at day 4 within a subset of activated cells. A similar behaviour was observed for *Tyk2* and *Tigit,* which encodes a receptor that is expressed by a subset of Tfh cells and was found to promote Tfh function (Godefroy et al., 2015). In contrast, expression of the Treg-associated gene *Ikzf4, Ly6c1,* which is expressed by effector T cells and, intriguingly, *Tbx21,* encoding the Th1 lineage-defining transcription factor Tbet become less variable between day 2 and day 4 during CD4^+^ T cell differentiation.

Comparing changes in mean expression and expression variability of *Cxcr5* and *Tbx21* transcripts across all time points shows that these lineage-defining genes exhibit opposite behaviours (**Fig. 4D**). While expression of both genes is up-regulated between days 2 and 4 in the lead-up to lineage commitment, *Cxcr5* increases and *Tbx21* decreases in variability. The fact that variability of *Tbx21* (Tbet) expression was highest on day 2 suggests that Tbet is up-regulated very early in differentiation, similar to *in vitro* Th1 induction (Szabo et al., 2000). Moreover, this suggests that Th1 fate decisions (for at least a subset of cells) may be made even earlier than the differentiation bifurcation point identified on day 4 by the original study (Lönnberg et al., 2017).

## Discussion

In recent years, the importance of modulating cell-to-cell transcriptional variation within cell populations for tissue function maintenance and development has become apparent (Bahar Halpern et al., 2015; Mojtahedi et al., 2016; Goolam et al., 2016). Here, we present a computational approach to robustly test changes in expression variability between cell populations using scRNAseq data. This extends the BASiCS framework by addressing experimental designs where spike-in sequences are not available and by incorporating an additional set of residual over-dispersion parameters *μ_i_*, which allow statistical testing of changes in variability that are not confounded by mean expression (Brennecke et al., 2013; Antolović et al., 2017). The latter is achieved by introducing a joint prior specification for the mean expression and over-dispersion parameters. We observe that this formulation stabilises posterior inference, particularly for small sample sizes. Furthermore, we show that our method successfully captures a mean versus over-dispersion trend across a variety of scRNAseq datasets.

Our method uncovers new insights into the extent and nature of variable gene expression in CD4^+^ T cell activation and differentiation. Firstly, we observe that during acute activation of naive T cells, the biosynthetic machinery is homogeneously up-regulated, while specific immune related genes become more heterogeneously up-regulated. In particular, increased variability in expression of the apoptosis-inducing Fas ligand (Strasser et al., 2009) and the inhibitory ligand PD-L1 (Chikuma, 2016) suggests a mechanism by which newly activated cells might suppress re-activation of effector cells, thereby dynamically modulating the population response to activation. Likewise, more variable expression of Smad3, which translates inhibitory *TGFβ* signals into transcriptional changes (Delisle et al., 2013), may indicate increased diversity in cellular responses to this signal. Increased variability in *Pou2f2* (Oct2) expression after activation suggests heterogeneous activities of the NF-ĸB and/or NFAT signalling cascades that control its expression (Mueller et al., 2013). Moreover, we detect up-regulated and more variable *Il2* expression, suggesting heterogeneous IL-2 protein expression known to enable T cell population responses (Fuhrmann et al., 2016). This emphasizes the importance of variability detection in scRNAseq data.

Finally, we studied changes in gene expression variability during CD4^+^ T cell differentiation towards a Th1 and Tfh cell state over a 7 day time course after *in vivo* malaria infection (Lönnberg et al., 2017). Our analysis provided several novel insights into this differentiation system. Firstly, we observe a tighter regulation in gene expression among genes that do not change in mean expression during differentiation at day 4, the bifurcation point where Th1 and Tfh differentiation separates. This decrease in variability on day 4 is potentially due to induction of a strong pan-lineage proliferation program. However, we observe that not all genes follow this trend and uncover four different patterns of variability changes. Secondly, we observe that several Tfh and Th1 lineage-associated genes change in expression variability between days 2 and 4. Importantly, we noted a decrease in variability for one key Th1 regulator, Tbx21 (encoding Tbet), which suggests that a subpopulation of cells had already committed to Th1 differentiation. Therefore, differentiation fate decisions may arise as early as day 2 in this system as reflected by high gene expression variability and in accordance with early commitment to Th1 and Tfh fates during viral infection (Choi et al., 2011). The diversity in differentiation state within this population of T cells can drive our differential variability results. Alternative analyses such as pseudotime inference used in Lönnberg et al. (2017) suggest that these differential variability results reflect a continuous differentiation process.

In sum, our model provides an important new tool for understanding the role of heterogeneity in gene expression during cell fate decisions. With the increasing use of scRNAseq to study this phenomenon, our and other related tools will become increasingly important.

## Methods

### The BASiCS framework

The proposed statistical model builds upon BASiCS (Vallejos et al., 2015, 2016) — an integrated Bayesian framework that infers technical noise in scRNAseq datasets and simultaneously performs data normalisation as well as selected supervised downstream analyses. BASiCS uses a hierarchical Poisson formulation to disentangle technical and biological sources of over-dispersion, corresponding to the excess of variability that is observed with respect to Poisson sampling noise. For this purpose, BASiCS uses a vertical integration approach that borrows information from synthetic spike-in RNA molecules (such as the set of 92 ERCC molecules developed by the External RNA Control Consortium, Jiang et al., 2011).

BASiCS summarizes the underlying distribution of gene expression across cells through two sets of gene-specific parameters. Firstly, mean expression parameters *μ_i_* capture average expression for each gene across all cells, matching what would be observed at a bulk level. Additionally, a second set of parameters *δ_i_* quantifies the biological over-dispersion that is observed for each gene after accounting for technical noise. Comparisons of these gene-specific parameters across populations can be used to identify statistically significant changes in gene expression at the mean and the variability level (**Fig. 1**). However, the well known confounding effect between mean and variability that typically arises in scRNAseq datasets (Brennecke et al., 2013) can preclude a meaningful interpretation of these results.

### Modelling the confounding between mean and over-dispersion with BASiCS

Here, we extend BASiCS to account for the confounding effect described above. For this purpose, we estimate the relationship between mean and over-dispersion parameters by introducing the following joint prior distribution for (*μ_i_,δ_i_*)’:

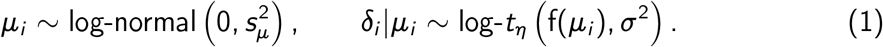

The latter is equivalent to the following non-linear regression model:

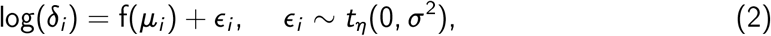

where f(*μ_i_*) represents the over-dispersion (on the log-scale) that is predicted by the global trend (across all genes) expressed at a given mean expression *μ_i_*. Therefore, *ϵ_i_* can be interpreted as a gene-specific *residual over-dispersion* parameter, capturing departures from the overall trend. If a gene exhibits a positive value for *ϵ_i_*, this indicates more variation than expected for genes with similar expression level. Similarly, negative values of *ϵ_i_* suggest less variation than expected.

A similar approach was introduced by DESeq2 (Love et al., 2014) in the context of bulk RNA sequencing. Whereas DESeq2 assumes normally distributed errors when estimating this trend, here we opt for a *t* distribution as it leads to inference that is more robust to the presence of outlier genes. Moreover, the parametric trend assumed by DESeq2 is replaced by a more flexible semi-parametric approach. This is defined by

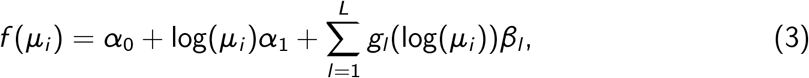

where *g*_1_(·), …, *g_L_*(·) represent a set of Gaussian radial basis function (GRBF) kernels and *α*_0_, *α*_1_, *β*_1_, …,*β_L_* are regression coefficients. As in Kapourani and Sanguinetti (2016), these are defined as:

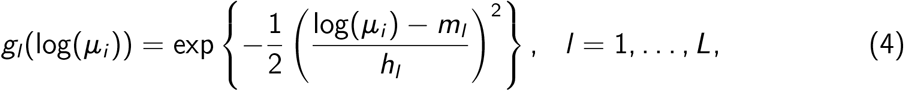

where *m_l_* and *h_l_* represent location and scale hyper-parameters for GRBF kernels.

In (3), the linear term captures the (typically negative) global correlation between *μ_i_* and *δ_i_*. Its addition also stabilises inference of GRBFs around mean expression values where only a handful of genes are observed.

### Implementation

Posterior inference is implemented through an Adaptive Metropolis within Gibbs sampler (Roberts and Rosenthal, 2009). The log-t prior in (1) is represented through a scale mixture of log-normal distributions as in Vallejos and Steel (2015). Moreover, the regression coefficients *β* = (*α*_0_, α_1_, *β*_1_,…, *β_L_*)’ are inferred by noting that (3) can be rewritten as a linear regression model using

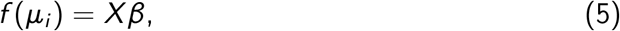

where *X* is a design matrix whose columns are a function of *μ_i_* (defined by the intercept, linear term and GRBFs in (3)). Finally, the Bayesian model is completed using a normal-inverse gamma prior on (*β, σ*^2^). More details about this implementation, including an explicit formula for *X* are described in **Supplementary Note S1**.

### Choice of hyper-parameters

In (3), the number of GRBFs and their associated hyper-parameters *m_l_, h_l_* are fixed *a priori.* As in Kapourani and Sanguinetti (2016), the location hyper-parameters *m_l_* are chosen to be evenly spaced across the range of mean expression values. Additionally, we set the scale hyper-parameters as *h_l_* = c × Δ*m*, where *c* is a fixed proportionality constant and Δm is the distance between consecutive values of *m_l_*. More details about these choices are provided in **Supplementary Note S2**. Default values for *L* and *c* are set as 10 and 1.2, respectively.

In principle, the degrees of freedom parameter *η* could be estimated within a Bayesian framework. Nonetheless, this is problematic (see Fernandez and Steel, 1999) and we observed that fixing this parameter *a priori* led to more stable results. Simulations across a range of values for this parameter showed that *η* = 5 provides a good compromise between shrinkage and flexibility (see **Supplementary Note S2**). We therefore set *η* = 5 as a default choice. While default values were set *a priori*, the model’s implementation also allows flexible adjustment of *L, c* and *η* by the user.

### Assessing changes in residual over-dispersion

We use a probabilistic approach to identify changes in gene expression between groups of cells. Let 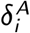 and 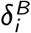 be the over-dispersion parameters associated to gene *i* in groups *A* and *B*. Following (2), the log_2_ fold change in over-dispersion between these groups can be decomposed as:

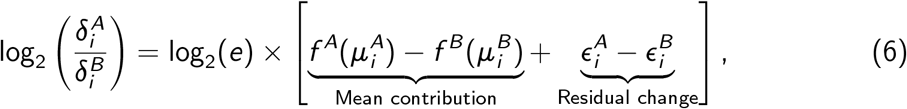

where the first term captures the over-dispersion change that can be attributed to differences between 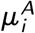 and 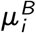. The second term in (6) represents the change in residual over-dispersion that is not confounded by mean expression. Based on this observation, statistically significant differences in residual over-dispersion will be identified for those genes where the tail posterior probability of observing a large difference between 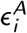 and 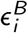 exceeds a certain threshold, i.e.

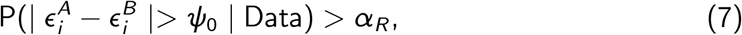

where *φ*_0_ > 0 defines a minimum tolerance threshold. As a default choice, we assume *φ*_0_ = log_2_(1.5)/ log_2_(*e*) ≈ 0.41 which translates into a 50% increase in over-dispersion. In the limiting case when *φ*_0_ = 0, the probability in (7) is equal to 1 regardless of the information contained in the data. Therefore, as in Bochkina and Richardson (2007), our decision rule is based on the maximum of the posterior probabilities associated to the one-sided hypotheses 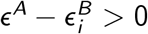 and 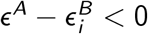, i.e.

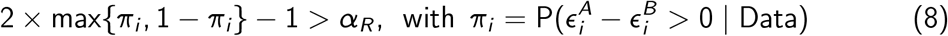

In both cases, the posterior probability threshold *α_R_* is chosen to control the expected false discovery rate (EFDR) (Newton et al., 2004). The default value for EFDR is set to 10%. As a default and to support interpretability of the results, we exclude genes that are not expressed in at least 2 cells per condition from differential variability testing.

## Data preparation

We employed a range of different datasets to test the proposed methodology. These datasets were selected to cover different experimental techniques (with and without unique molecular identifiers, UMI) and to encompass a variety of cell populations: CD4^+^ T cells (Martinez-Jimenez et al., 2017), microglia cells and CA1 pyramidal neurons (Zeisel et al., 2015), cells from the differentiating *Dictyostelium* (Antolović et al., 2017) and an experimental control to study technical noise (Grün et al., 2014). In all cases, the analysis is based on raw data obtained from publicly available links. The gene and cell filter quality control criteria applied to each dataset are described in **Supplementary Note ??** and key features of each dataset can be found in **Table S1**.

## Software availability

Software (R) is freely available at github.com/catavallejos/BASiCS/tree/Devel and will be released as part of Bioconductor 3.7.

R scripts for data preparation and analysis are available at github.com/MarioniLab/RegressionBASiCS2017.

## Acknowledgements

NE was funded by the European Molecular Biology Laboratory (EMBL) international PhD programme. ACR was funded by the MRC Skills Development Fellowship (MR/P014178/1). SR was funded by MRC grant MC_UP_0801/1. JCM was funded by core support of Cancer Research UK and EMBL. CAV was funded by The Alan Turing Institute, EPSRC grant EP/N510129/1.

## Author Contributions

N.E., J.C.M., and C.A.V. designed the study; N.E. and C.A.V build the model and performed computational analyses; N.E. and A.C.R performed biological data interpretation; S.R. provided technical assistance; and N.E., A.C.R., J.C.M., and C.A.V. wrote the manuscript. All authors commented on and approved the manuscript.

